# scSpark: an AI-assisted cloud platform for traceable interpretation of single-cell transcriptomic results

**DOI:** 10.64898/2026.07.03.736259

**Authors:** Jinbo Zhang, Zhengxi Liu, Zhongqi Pu, Da Mi

## Abstract

Single-cell RNA sequencing now routinely produces detailed maps of cell types and states, but interpreting a finished project remains harder than it should be. Once the analysis is done, the results are usually handed over as static reports, figure panels and supplementary tables. A biologist who later wants to revisit an annotation, recompute a cell-type proportion or check whether a pathway is specific to one group typically has to return to a bioinformatician rather than explore the data directly. We developed scSpark to close this gap. The platform takes the completed outputs of a single-cell project: cell annotations, embeddings, differential-expression tables, trajectories, cell-cell communication networks and enrichment results—and serves them through a web browser as an interactive workspace. Heavy computation stays upstream: scSpark indexes the precomputed objects under a single project structure and exposes them through six modules for cell annotation, differential analysis, trajectory exploration, cell-cell communication, functional interpretation and AI-assisted result interrogation. Every action in these modules, from a query to a label change, an export or an AI-generated summary, is linked to a specific project version, data object, parameter set and output file, so that any conclusion can be traced back to the evidence behind it. We illustrate the platform by reworking a published periodontitis dataset through this interface. scSpark does not replace upstream pipelines or expert judgement; it is a layer that makes their results easier to inspect, revise and reuse, and that turns a single-cell project from a one-off report into an interpretation others can follow and check.

**Significance Statement:** Single-cell studies produce increasingly intricate maps of tissues, but those maps are hard to interrogate once they have been written up as static reports. scSpark tackles this post-analysis bottleneck by holding a project’s annotations, marker evidence, differential results, pathways, communication networks, AI-generated summaries and publication-ready figures together in one workspace, where each item is linked to the data and settings that produced it. The platform is built to support expert decisions rather than to make them: its aim is to let researchers check, revise and reuse a result, and to see exactly how it was reached.

## Introduction

Single-cell profiling has become a standard way to study complex tissues. The earliest single-cell mRNA-sequencing work showed that a whole transcriptome could be read out from one cell.^1^ Droplet methods soon made it routine to profile thousands of cells at once,^2^ and large consortia, notably the Human Cell Atlas, turned single-cell genomics into a framework for cataloging human cell types, states and tissue programs.^3^ Spatial transcriptomics added a further dimension, tying expression back to tissue architecture so that molecular states can be read in their anatomical context.^4^ Between them, these technologies have pushed many questions past bulk-tissue comparisons into cellular composition, state transitions, tissue niches and intercellular signaling.

Getting value out of these assays takes a long chain of computational and judgment calls. A typical project runs through quality control, normalization, feature selection, dimensionality reduction, batch correction, clustering, annotation, differential expression, pathway analysis, trajectory inference and cell-cell communication analysis,^5^ and widely used frameworks such as Seurat and Scanpy have made most of these steps reproducible and accessible to non-specialists.^6-8^ Standardization, however, has not removed the need for expert review. Where clusters begin and end, whether a marker is specific, how finely to annotate, and whether a population is a stable cell type, a transient state, an artifact or a disease-associated program are questions that usually demand repeated inspection of the evidence.^9^ A biological conclusion is seldom settled by one plot or one test; it is built up from agreement across annotations, marker genes, sample composition, differential expression, pathway activity and prior knowledge.

How projects are delivered makes this harder. In day-to-day research and in service work, the analyst usually hands over static reports, image panels and supplementary tables once the upstream analysis is finished. Those files record an answer, but they do little to support the back-and-forth through which a biologist actually refines a hypothesis: comparing alternative labels for a cluster, checking the marker evidence behind it, asking whether a pathway is specific to one condition, testing whether a ligand-receptor signal comes from only a handful of cells, or re-exporting a figure for a paper. Interactive viewers have helped. ShinyCell turns a dataset into a shareable interface,^10^ and CZ CELLxGENE Discover scales exploration to large curated collections.^11^ But interpreting a whole project takes more than a viewer. It also needs versioned data objects, traceable parameters, editable annotations, standardized exports, and a record of how each figure and table came to be.

Interpretation also leans on structured biological knowledge. Gene Ontology and KEGG supply the controlled vocabularies and pathway maps that put gene-level results in context,^12,13^ while gene-set enrichment analysis and GSVA read interpretation at the pathway level from ranked genes or expression profiles.^14,15^ For intercellular signaling, frameworks such as CellChat and CellPhoneDB infer candidate ligand-receptor interactions from the same single-cell data.^16,17^ Each of these is most informative not on its own but as one strand of evidence in a shared workspace. A candidate disease-associated population, for instance, is far more convincing when its markers, its shift in abundance, its differential genes, its enriched pathways, its position along a trajectory and its communication patterns can all be examined side by side rather than scattered across separate files.

Foundation models and language-model systems are beginning to make this interpretive layer more interactive. scGPT, for example, shows that large single-cell datasets can be represented in a form that supports annotation, integration and perturbation analysis,^18^ and multi-agent language-model systems have started to annotate cells while exposing their reasoning and flagging quality.^19^ This is promising, but it sets a clear requirement for scientific software: any automated interpretation has to stay tied to evidence that can be audited. A generated summary helps only if the user can follow it back to the data object, the module, the parameters and the source tables behind it; otherwise the convenience comes at the cost of reproducibility, and the conclusions are harder to defend.

This is the gap scSpark is built to fill. It is a cloud platform that takes the finished outputs of a single-cell project and organizes them into an interactive, traceable workspace for downstream interpretation, without replacing the upstream pipeline. Precomputed objects, metadata, annotations, marker-gene tables, differential-expression results, enrichment analyses, trajectory states, ligand-receptor networks and user-defined gene-set scores are all indexed so that they can be queried and exported. In one place, a user can check marker support, relabel cells, compare cell fractions between groups, pull out condition-specific genes, query GO, KEGG, GSEA and GSVA results, inspect inferred communication, and produce publication-ready figures and tables without writing code. Every one of these actions is tied to a specific project version, data object, module, parameter set and downloadable file. By replacing one-off report delivery with a workspace that can be revisited, scSpark makes single-cell results easier to inspect, reuse and turn into testable hypotheses.

## Results

### scSpark implements a project-oriented cloud workflow

scSpark is implemented as a server-side web application that requires no local installation of R, Python or associated single-cell analysis environments. Users interact with the platform through a standard web browser. After authentication, the server resolves the project and batch associated with the user, reads the corresponding configuration files and dynamically loads the modules enabled for that project. This design separates project access, data storage and analysis modules from the client interface, making the workspace easier to maintain across projects (Fig. 1).

**Fig. 1.**
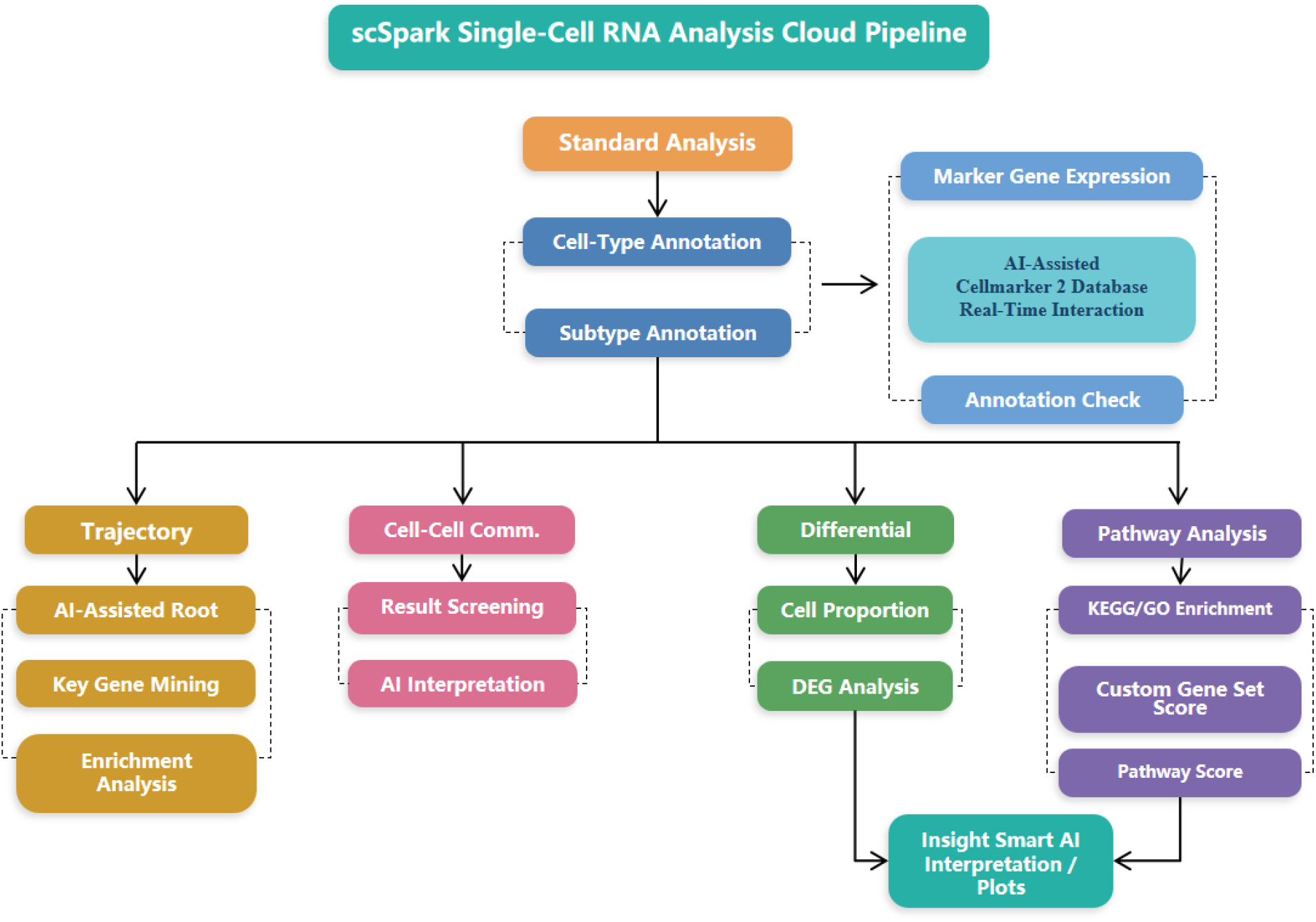
Overview of the scSpark workflow. scSpark separates standardized upstream analysis from interactive downstream interpretation. After a user signs in, the platform resolves the project and analysis batch, loads the modules enabled in the project configuration, and serves precomputed results through six browser modules (cell annotation, differential analysis, trajectory exploration, cell-cell communication, functional pathway analysis and AI-assisted Insight). Tables, figures and AI-generated code can be downloaded, and each output is recorded with its data object, parameter settings and project version to support reproducibility.

The interface follows a consistent two-panel layout. A left-hand panel manages authentication state, project and batch selection and module navigation, while the right-hand panel renders the selected module. Each project is organized under a project identifier and a batch identifier. The corresponding directory contains project-level statistics, sample metadata, a configuration file, dedicated subdirectories for each class of analysis result and a temporary directory for on-demand outputs (Table 2). Because module availability, file paths and default parameters are declared in a standardized configuration file and verified by a file-validation step, scSpark can be adapted to new projects through configuration rather than code modification.

### Six modules support post-analysis interpretation

scSpark integrates six modules within a single interface: cell annotation, differential analysis, trajectory exploration, cell-cell communication, functional pathway analysis and AI-assisted Insight (Table 1 and Fig. 2). Together, these modules cover the tasks most frequently required after upstream single-cell processing: inspection of cell-population annotations, screening of differentially expressed genes, browsing of pseudotime structures, interrogation of CellChat-based communication results, GO and KEGG enrichment, gene-set scoring, transcription-factor or pathway scoring and AI-assisted interpretation of selected result files.^12,13,16,21-23^

**Table 1.** Summary of scSpark analytical modules.

| Module | Purpose | Main inputs | Main outputs | Runtime mode |
| --- | --- | --- | --- | --- |
| Cell annotation | Annotation review and label refinement | Annotation file, marker tables, UMAP coordinates | Annotated UMAP, annotation table, marker evidence | Precomputed + review |
| Differential analysis | Cell-fraction comparison and DEG screening | Annotation, DEG tables, expression matrix | Proportion plots, filtered gene tables, set intersections | Precomputed + on-demand |
| Trajectory exploration | Pseudotime and branch inspection | Monocle-compatible trajectory objects | Pseudotime embeddings, branch structure, dynamic genes | Precomputed |
| Cell-cell communication | Ligand-receptor and pathway-level interaction browsing | CellChat object and interaction tables | Communication networks, pathway plots, ligand-receptor tables | Precomputed |
| Functional pathway | GO/KEGG enrichment and gene-set scoring | Gene lists, signature files, scoring matrices | Enrichment tables, pathway scores, downloadable outputs | On-demand |
| AI-assisted Insight | Result interpretation and figure-code generation | Selected result file | Interpretive summary, visualization suggestions, editable R code | On-demand model call |

**Table 2.** Standardized project directory contents.

| Path or file | Contents | Use in scSpark |
| --- | --- | --- |
| project_id/batch_id/project_stat.tsv | Project-level summary statistics | Overview page and quality summary |
| project_id/batch_id/sample_info.tsv | Per-sample metadata | Group selection and sample-level summaries |
| project_id/batch_id/project_conf.json | Module flags, result-file paths and defaults | Configuration-driven module loading |
| annotation/ | Annotation files, marker evidence and UMAP coordinates | Cell annotation module |
| diff/ | Precomputed DEG tables and grouping definitions | Differential analysis module |
| trajectory/ | Monocle-compatible objects and trajectory outputs | Trajectory exploration module |
| celltalk/ | CellChat objects and interaction outputs | Cell-cell communication module |
| enrich/ | Enrichment results, signature files and scoring outputs | Functional pathway module |
| insight/ | Insight-related result files and templates | AI-assisted interpretation |
| tmp/ | Temporary on-demand outputs | Downloadable intermediate results |

**Fig. 2.**
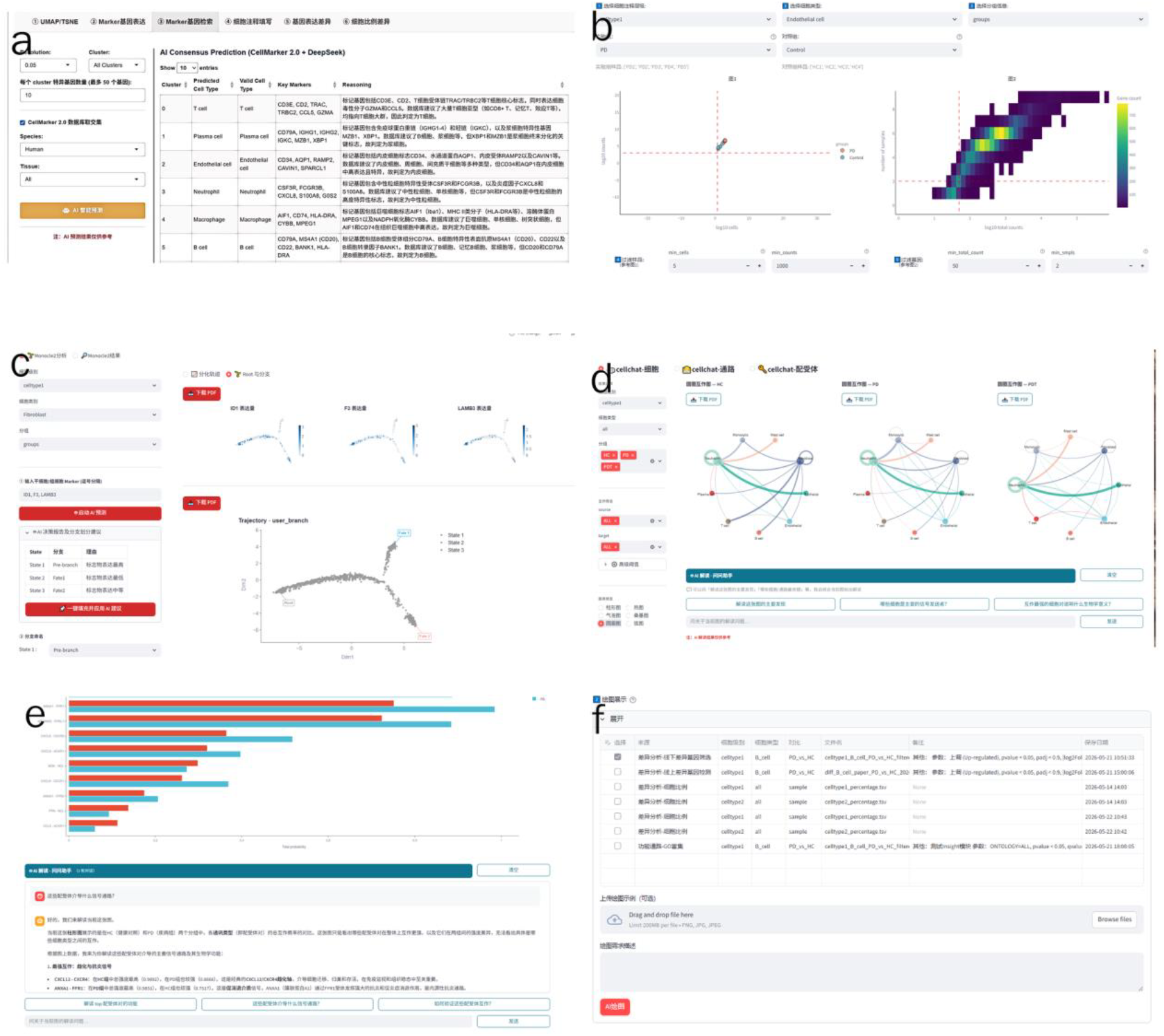
Interactive analysis modules of scSpark. Browser views of the six modules: (a) AI-assisted cell annotation, combining a curated marker reference with a large language model; (b) differential analysis with on-demand group and parameter selection; (c) trajectory exploration based on Monocle pseudotime; (d) cell-cell communication based on CellChat; (e) functional pathway analysis with enrichment results and an interpretation panel; and (f) the Insight module, which links a selected result file to a summary and editable plotting code.

The platform is designed for immediate interaction rather than exhaustive reparameterization of upstream methods. Each module exposes only the core options needed for exploration, such as samples, groups, cell types, genes, pathways or result files. Outputs are rendered directly in the browser as tables, figures and status messages. This design lowers the operational barrier for users without programming experience while keeping each result connected to its underlying file and parameter state.

### A public periodontitis dataset illustrates project-level reuse

To illustrate typical use, we applied scSpark to a published single-cell RNA-sequencing study of the osteoimmunology microenvironment in periodontitis.^24^ The dataset contains 51,248 cells from healthy controls, patients with severe chronic periodontitis and patients sampled within one month after initial periodontal therapy. A user first inspects sample composition and cell numbers on the project overview page, then enters the annotation module to review annotation status and the uniform manifold approximation and projection distribution of annotated populations.

Using the annotation module together with a manually curated marker reference, including CellMarker 2.0,^21^ scSpark recovered ten major cell populations: T cells, plasma cells, endothelial cells, neutrophils, monocytic cells, B cells, epithelial cells, fibroblasts, mast cells and pericytes (Fig. 3 and Table 3). These populations cover the major fibroblast, monocytic, endothelial, T-cell and B-cell compartments described in the original cell atlas. The purpose of this demonstration was not to reproduce every analysis in the original study; rather, it shows how a completed single-cell project can be reorganized into an interactive workspace in which annotation, differential analysis, functional interpretation and trajectory evidence are inspected through a single interface.

**Table 3.** Representative cell-type annotations recovered in the periodontitis demonstration project.

| Cluster | Predicted cell type | Representative markers |
| --- | --- | --- |
| 0 | T cell | CCL5, CD2, CD3E, CD3G, TRAC, TRBC1, TRBC2, GZMA, IL7R |
| 1 | Plasma cell | CD79A, IGHG1, IGKC, IGLC2, MZB1, XBP1, DERL3 |
| 2 | Endothelial cell | CD34, PECAM1, VIM, RAMP2, AQP1, SPARC |
| 3 | Neutrophil | S100A8, S100A9, IL1B, CXCL8, CXCR2, CSF3R, FCGR3B |
| 4 | Monocytic | CD74, HLA-DRA, HLA-DPA1, HLA-DPB1, AIF1, CST3, FCER1G, MPEG1 |
| 5 | B cell | BANK1, CD22, CD79A, MS4A1, HLA-DRA |
| 6 | Epithelial cell | KRT5, KRT6A, S100A8, S100A9, SPRR2A, TACSTD2 |
| 7 | Fibroblast | COL1A1, COL1A2, COL3A1, DCN, LUM, SPARC, VCAN |
| 8 | Mast cell | TPSAB1, TPSB2, CPA3, KIT, HDC, MS4A2 |
| 9 | Pericyte | CALD1, MYLK, NOTCH3, RGS5, MCAM, SPARC |

**Fig. 3.**
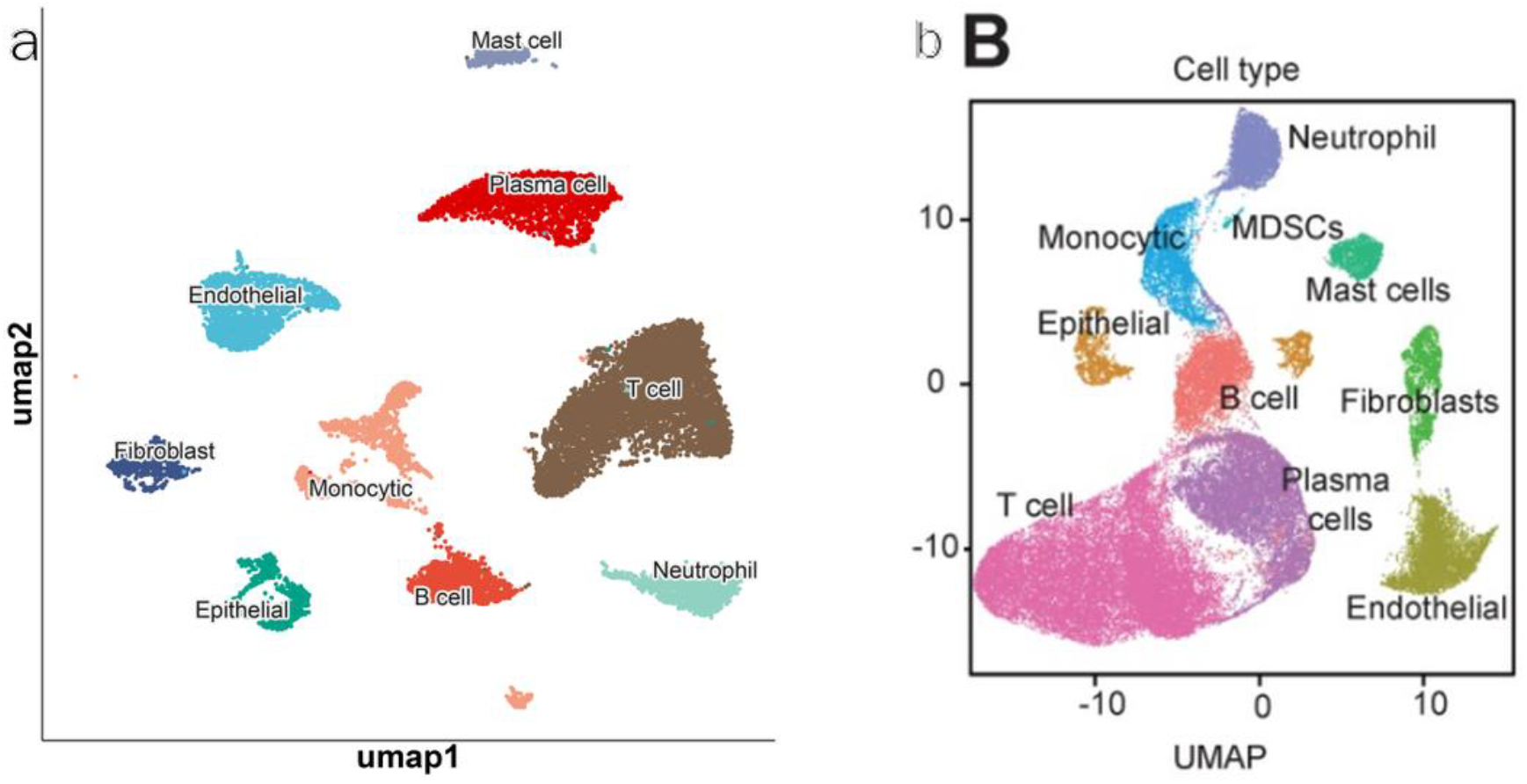
Cell-type annotation in the periodontitis demonstration. (a) UMAP of cell-type annotations produced by scSpark. (b) Corresponding annotations reported in the original study. The platform recovers the major cell populations of the published atlas through its annotation module.

Within the same workspace, users can compare cell-type proportions across clinical groups, screen differentially expressed genes, carry candidate gene sets into GO and KEGG enrichment, inspect pathway scores and examine precomputed trajectory or communication results (Fig. 4). Selected tables can then be passed to the Insight module for interpretation and figure planning. This workflow keeps numerical outputs, biological interpretation and figure export within one project context, reducing the need to move repeatedly among disconnected files.

**Fig. 4.**
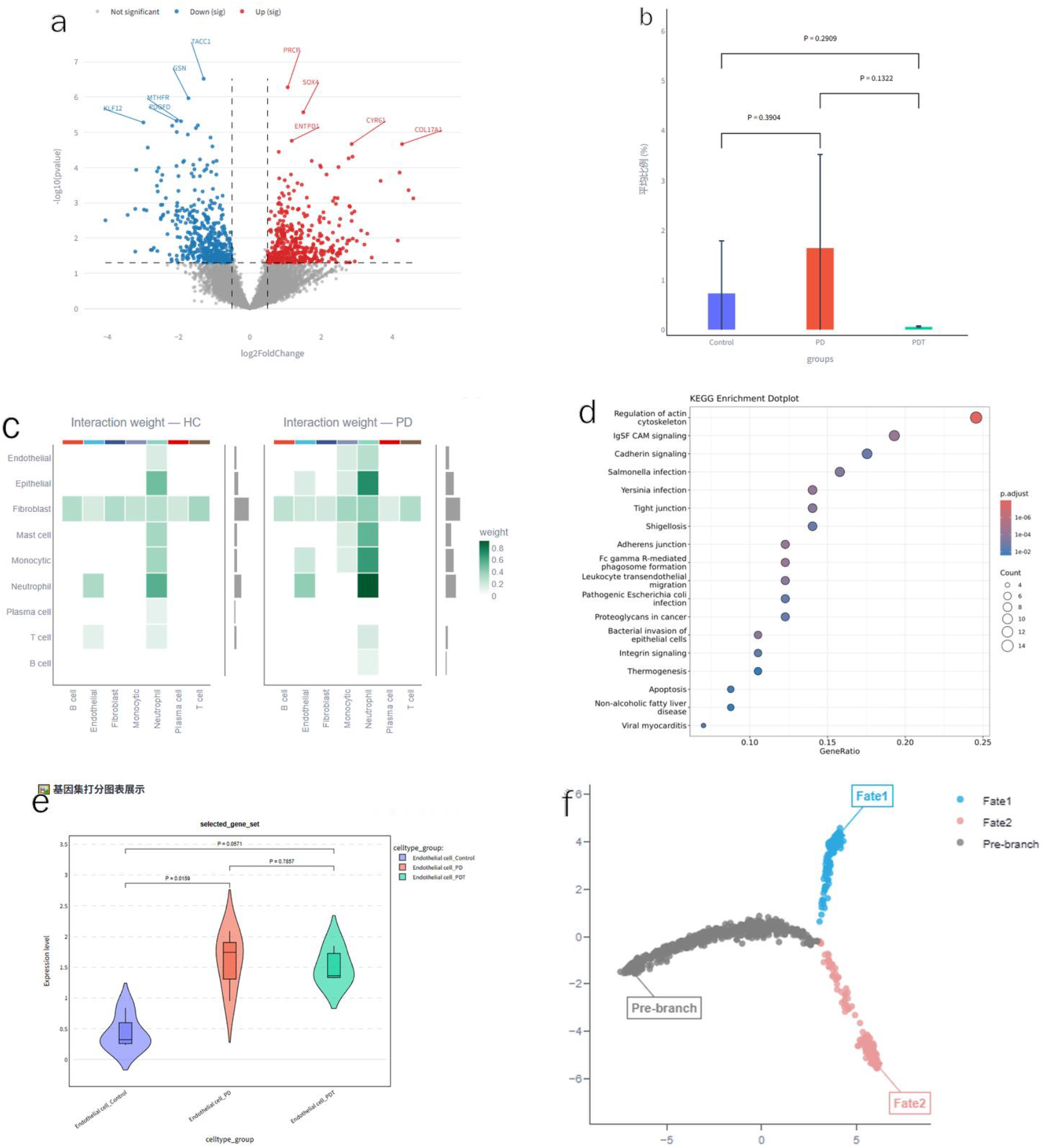
Downstream interpretation in the periodontitis demonstration. (a) Differential expression between clinical groups, shown as a volcano plot. (b) Cell-type proportions across healthy-control, periodontitis and post-therapy groups. (c) Cell-cell communication strength in healthy-control and periodontitis samples. (d) KEGG enrichment of differentially expressed genes. (e) Gene-set and pathway activity scoring across groups. (f) Pseudotime trajectory with inferred branches. All panels were produced with scSpark’s standard modules.

### Traceable AI-assisted interpretation links results to decisions

scSpark emphasizes outputs that users can retain, verify and reuse. Tables generated during browsing or on-demand analysis, including differential-gene tables, annotation files, enrichment results, ligand-receptor interaction tables, gene-set scoring outputs and AI-generated code, are made directly downloadable. Figures can be saved through standard browser actions or regenerated from the underlying result files using the same parameter selections.

The Insight module connects selected result files to biological interpretation. Given a user-selected table or result file, the module returns a concise interpretive summary, suggests suitable visualization types and generates editable R code for downstream plotting. The module is positioned as an interpretation aid rather than a discovery system. Its outputs require human verification, it does not draw on data sources beyond the project files unless explicitly configured and it falls back to direct file inspection whenever a file cannot be interpreted. By coupling result files with interpretation and visualization scaffolds, the Insight module reduces the operational distance between numerical outputs and biological reasoning while preserving user oversight.

## Discussion

The problem scSpark addresses is not a shortage of computational tools but the distance between a finished analysis and an interpretation a biologist can defend. Quality control, normalization, batch correction, dimensionality reduction, clustering, trajectory inference and differential expression can all be run at scale with standard workflows.^5-8^ What still rests on expert judgment are the steps that decide a study: whether a cluster is a real cell state, whether a marker set holds up, whether a pathway result is mechanistically plausible, and whether a figure can carry a claim in a manuscript. Automating the pipeline speeds up the computation but does not, on its own, make any of these calls easier. We designed scSpark for this later stage, where an investigator keeps moving between the data, the annotations, the literature and a hypothesis that is still taking shape.

That focus sets scSpark apart from existing analysis and visualization resources. Seurat and Scanpy are flexible environments for running the analysis itself,^6-8^ and ShinyCell and CELLxGENE have made the resulting datasets much easier to share and explore.^10,11^ Both have been important for opening up single-cell data. scSpark works downstream of them, shifting the emphasis from browsing data to supporting project-level decisions: instead of leaving plots, tables, annotations, enrichment results and communication networks as separate files, it ties them together as parts of one workspace. That matters most in collaborative and service projects, where a dataset is often revisited as the biological question changes.

It also implies that reproducibility should reach past the upstream pipeline. In single-cell work, many of the decisions that matter happen after clustering: labels are revised, marker lists are filtered, the resolution is reconsidered, candidate states are split or merged, and particular pathway results are chosen to interpret. These choices are rarely recorded as carefully as the preprocessing is. scSpark’s traceability layer ties each output back to the data object, parameters, annotation version and export that produced it, in keeping with the FAIR principles, under which data, algorithms, tools and workflows should be findable, accessible, interoperable and reusable.^20^ For a single-cell project, it means a final figure is never separated from the analytical choices behind it.

Adding AI assistance changes what a platform is for. Foundation models and language-model systems have shown that large single-cell datasets and the biological literature can together support annotation, representation learning and the generation of interpretable results.^18,19^ The danger is familiar: a fluent explanation can read as convincing even when the evidence under it is thin. For that reason scSpark does not hand interpretation over to an unconstrained model. Its AI functions sit inside bounded modules that summarize marker evidence, rank candidate mechanisms, suggest lines of interpretation and help lay out figures and tables, always with the analyst in the loop. The dependable near-term role for AI here is a modest one: helping researchers see more of their data, weigh more alternatives, and record why they accepted a given interpretation.

How useful the system is depends on the evidence it can bring together. scSpark holds annotation evidence, differences in cell composition, differential expression, trajectory analysis, ligand-receptor inference and pathway-level interpretation in one place. None of these is a conclusion on its own; each is a different view of the same biological system. A disease-associated population becomes credible when its abundance shifts with condition, its markers hang together, its trajectory position makes sense, and its inferred signaling points to a microenvironmental mechanism that can be tested. This is how single-cell studies are read in practice, with ontology, pathway knowledge, enrichment and communication inference serving as supporting layers rather than endpoints in themselves.^12-17^

The platform has clear limits. It cannot rescue a study from poor sample design, too few recovered cells, low sequencing quality, unmodeled batch effects or unsuitable upstream filtering. Annotation stays context dependent: reference atlases and marker databases help, but they cannot fully resolve rare, transitional, activated or disease-specific states in tissues where reference knowledge is thin. Enrichment and ligand-receptor analyses are hypothesis-generating, surfacing coherent signals and candidate mechanisms without proving causality on their own. And the AI summaries have to stay auditable, with every generated interpretation traceable to marker genes, statistics, pathway terms or evidence the user supplied.

Several directions follow from this. Because scSpark is an interpretation platform rather than a single algorithm, the usual accuracy metrics are not enough to evaluate it; more telling benchmarks would ask whether it improves annotation consistency, shortens turnaround, makes figures more reproducible, preserves a decision history, and lets independent users reach the same biological conclusions from the same project. As studies increasingly combine RNA with chromatin accessibility, surface protein, spatial position, immune-receptor sequence and perturbation readouts, the platform should treat these as one project object rather than separate streams. And for clinical data, private deployment and access control will matter, since the convenience of the cloud has to be weighed against data governance and institutional rules.

In the end, scSpark treats single-cell analysis as an iterative decision process rather than a one-time computation. It is not meant to replace established methods, but to make their results easier to inspect, revise, interpret and document. By bringing standard analyses, AI assistance, expert review and reproducible export into a single workspace, it turns a static report into something a group can keep working in. That seems a practical direction for single-cell omics: keep the computation rigorous, keep interpretation in human hands, and make the path from data to manuscript-ready evidence transparent enough to check, share and reuse.

## Methods

### Platform architecture and deployment

scSpark is implemented as a server-side web application accessed through a standard web browser. Application logic, data resolution and file validation run on the server, while the client renders interface elements and analysis outputs. The interface adopts a two-panel layout, with authentication, project and batch selection and module navigation on the left, and module-specific results on the right. This design removes the need for users to maintain local R, Python or single-cell analysis environments.

### Project-based authentication and access control

Access is organized at the level of individual projects. Following authentication, the platform resolves the project and batch associated with the user and restricts directory access to the corresponding server-side paths. Directory resolution is performed on the server, and absolute project paths are not exposed to the client.

### Project directory and configuration schema

Each project is stored under a standardized directory keyed by project_id and batch_id (Table 2). The directory contains project-level summary statistics, per-sample metadata and a configuration file that declares which modules are enabled, the paths to relevant result files and default parameter values. Dedicated subdirectories hold annotation, differential, trajectory, communication, enrichment and Insight outputs, together with a temporary directory for on-demand results. At load time, the configuration is parsed and a file-validation step confirms the presence of declared files, allowing new projects to be deployed through configuration.

### Cell annotation module

The annotation module reads precomputed annotation files stored as Seurat-compatible objects and renders annotation status, downloadable annotation tables and UMAP visualizations of annotated cell populations.^6,8^ UMAP coordinates are computed during project preparation and read from the stored object. AI-assisted annotation suggestions are derived from project marker tables, a curated marker reference such as CellMarker 2.0,^21^ and an optional language-model prompt. Suggested labels are presented for user review before adoption.

### Differential analysis module

The differential-analysis module supports comparison of cell-type proportions between groups, filtering of precomputed differential-gene tables and on-demand screening for user-selected groups and cell types. Where appropriate, expression values can be aggregated into per-sample pseudo-bulk profiles before testing. Set-intersection analysis across comparisons is provided to identify shared and unique genes. Inputs include annotation results, differential-gene tables and expression matrices; outputs include proportion plots, filtered gene tables and Venn-style summaries.

### Trajectory exploration module

The trajectory module renders precomputed pseudotime results generated during project preparation, including Monocle-based outputs where available.^22^ Users can browse pseudotime-ordered embeddings, branch structures and dynamic gene-expression profiles along the trajectory. Differentiation roots can be specified manually or guided by marker evidence, and gene dynamics can be linked to differential-expression and enrichment results.

### Cell-cell communication module

The communication module displays precomputed CellChat results at the levels of cells, signaling pathways and ligand-receptor pairs.^16^ Users can select groups for comparison, display multiple conditions simultaneously and restrict inspection to specified sender and receiver cell types. Outputs include communication networks, pathway-level plots and ligand-receptor tables.

### Functional pathway module

The functional-pathway module supports GO and KEGG over-representation analysis on user-selected gene lists, as well as gene-set and pathway activity scoring using predefined signature files.^12-15,23^ Enrichment and scoring are executed on demand within constrained parameter ranges exposed by the interface. Outputs include enrichment tables and scoring matrices that can be inspected and downloaded.

### AI-assisted Insight module

The Insight module connects result files to biological interpretation. For a user-selected result file, the module returns a natural-language summary, suggests appropriate visualization types and generates editable R code for figure construction. Generated code and summaries are presented for inspection and can be copied, edited and downloaded. The module does not access data beyond the project result files unless this behavior is explicitly configured.

### Runtime strategy and reproducibility logging

Interactive responsiveness is achieved by separating offline precomputation from online execution. Computationally intensive steps, including cell annotation, communication inference and pseudotime estimation, are precomputed during project preparation and stored as serialized objects. On-demand computation at browse time is limited to lightweight operations, including proportion comparison, gene-table filtering, set intersection and enrichment analysis. Each query, parameter selection, output file and exported result is associated with the project and batch context, supporting regeneration of displayed results from recorded objects and settings.

## Data availability

No new sequencing data were generated in this study. The representative scRNA-seq dataset used for demonstration derives from the periodontitis single-cell study of Chen et al.^24^ and is available from the Gene Expression Omnibus under accession GSE171213.

## Code and software availability

scSpark is a web-based platform. A demonstration instance, together with access details for the example periodontitis project, is available at [URL]. [State the access and licensing terms: if the source code is openly available, give the repository link and license; if the platform is proprietary or access-controlled, state how qualified users can obtain access.] Software dependencies and their versions used for project preparation and deployment are listed in Supplementary Table 2.

## Acknowledgements

We thank all the users who provided feedback during platform testing and interface refinement.

## Author contributions

J.B.Z. conceived and led the development of scSpark, including overall design, functional planning, interface layout and system implementation. Z.Q.P. and Z.X.L. provided code support and optimized the user interface. All authors reviewed and approved the manuscript.

## Supplementary information

Supplementary Fig. 1 | System architecture and project directory schema. Server-side application architecture, project-based authentication and standardized project_id/batch_id directory organization.

Supplementary Fig. 2 | Representative interface views of the six analysis modules.

Supplementary Fig. 3 | Insight module input-output examples for supported result-file types.

Supplementary Table 1 | Schema for downloadable result files across modules.

Supplementary Table 2 | Software dependencies and version numbers used for project preparation and deployment.

Supplementary Note 1 | Annotated project configuration template.

Supplementary Note 2 | Demonstration project access instructions and credential policy.

